# The influence of marker number and sequencing depth on the ability to identify mismatch repair deficient tumours

**DOI:** 10.1101/2023.07.18.546679

**Authors:** Joseph Law, Richard Gallon, Ethan Teare, Ivan Santibanez Koref, Rachel Phelps, John Burn, Michael Jackson, Mauro Santibanez Koref

## Abstract

Analysis of somatic mutation patterns is widely used to infer exposure to exogenous and endogenous mutagenic influences. This raises the question of the amount of sequence data required to detect factors of interest. A common use of mutation pattern analysis is the identification of increased microsatellite instability to uncover mismatch repair (MMR) defects in tumours and normal tissues. Here we explore the effects of sequencing depth and the number of loci analysed on the ability to detect MMR deficiency using artificial neural networks and publicly available amplicon sequencing data from colorectal tumours on 24 short quasi monomorphic microsatellites (up to 12 bp in length, PMID 31471937) split in a training (99 samples) and a test set (95 samples). We show that, at a sequencing depth of 200, pairs mononucleotide repeats can achieve discrimination between MMR proficient and deficient colorectal tumours similar to that obtained with the full 24 marker panel, with accuracies above 97% and ROC AUCs in excess of 99% in the test set. Our results indicate that for short monomorphic microsatellites considering the length distribution of the different alleles at each locus, representing these distributions as two-dimensional structures and including convolutional layers in the network can facilitate discrimination between MMR deficient and proficient tumour material. They also indicate that, despite the limitations imposed by amplification, sequencing accuracy and the limited divergence time between the sequences from one locus, high depth sequencing can be used to identify MMR deficiency from a limited number of loci. However, they also suggest that, for a fixed total number of reads per sample, increasing sensitivity by increasing the number of targets is more efficient than by increasing per target sequencing depth. These results are of interest for screening large numbers of samples and for assessing the impact MMR deficiency in different areas of the genome.

## 2. Introduction

The DNA in the cells of the body is constantly exposed to a variety of endogenous and environmental factors that can induce sequence changes and, if left unrepaired, to variation between the genomes of the cells in the body. Different factors lead to mutations with different characteristics and patterns of somatic mutations have been widely used to infer exposure to exogenous and endogenous mutagenic influences ^1,2^. Perhaps one of the most common uses of mutation pattern analysis is the identification of tumours lacking a functional mismatch repair (MMR) system, one of the mechanisms responsible for correcting mutations ^3,4^. Tumours lacking functional mismatch repair are particularly susceptible to specific therapies like immune checkpoint blockage inhibition and resistant against others such as treatment with alkylating agents ^5^. The MMR system includes a number of genes whose function can be impaired by mutations or epigenetic changes such as methylation of promoter regions ^4-7^.

The ability to identify a mutagen based on the characteristics of the mutations it has caused is influenced by several factors including number of mutations generated and the uniqueness of the mutation pattern induced. In general, the larger the number of mutations analysed the easier the identification of the contributing factors will be. This number will depend on the amount of DNA screened in the search for mutations and reflects the number of loci analysed and the number of genomes from different cells represented in the sequencing results. Each region of the genome may be sequenced more than once, and these sequences may be derived from different cells. Here we refer to the number of times a particular region of the genome is represented in the sequencing results as the sequencing depth. Low sequencing depth means that at each position of the genome information from only a few cells is available, while a higher sequencing depth allows to assess the sequence of a particular region for a larger number of cells.

With the development of cheap whole genome sequencing methods, the analysis of large regions of the genome at low or moderate depths has become increasingly popular. An alternative is to sequence fewer loci but at higher depths. A high depth approach is particularly appealing if mutations are concentrated in well-defined regions of the human genome.

Mismatch repair deficiency is characterised by an increase of mutations of microsatellites. These are regions of the genomes consisting of tandem repetition of short (1 to 6 nucleotide long) DNA motifs. Mononucleotide repeats consisting of repetitions of the nucleotide adenine are commonly used to detect mismatch repair deficiency ^8,9^. Mismatch repair deficiency tends to induce mutations consisting of deletions or insertion of repeat units, leading to a change in the length of the microsatellites. Since microsatellites are prone to mutation both *in vivo* and *in vitro*, inferring mismatch repair deficiency from the frequency of novel microsatellites alleles involves separating the signal, i.e., the increase mutation rate due to the deficiency, from the noise due to the background mutation rate and to technical artifacts. Microsatellites used to track features of interest such as an increased mutation rate are often called microsatellite markers (or simply markers when the nature of the sequence element is clear from the context). We will call a deviation from the (putative) germline sequence a variant. We will refer to each single sequence obtained in a sequencing experiment as a read. In general, each read reflects the sequence of a single molecule of input material, but depending on the sequencing technology used, two reads may represent the sequence of the same input molecule.

Here we explore the relationship between sequencing depth and number of loci used in the analysis on the ability to identify mutagenic factors using mismatch repair proficient and deficient tumours as a model. Figure 1 illustrates some of the issues associated with increasing sequencing depth while maintaining the total number of reads. Issues arising from technical artifacts are disregarded in the figure.

**Figure 1.**
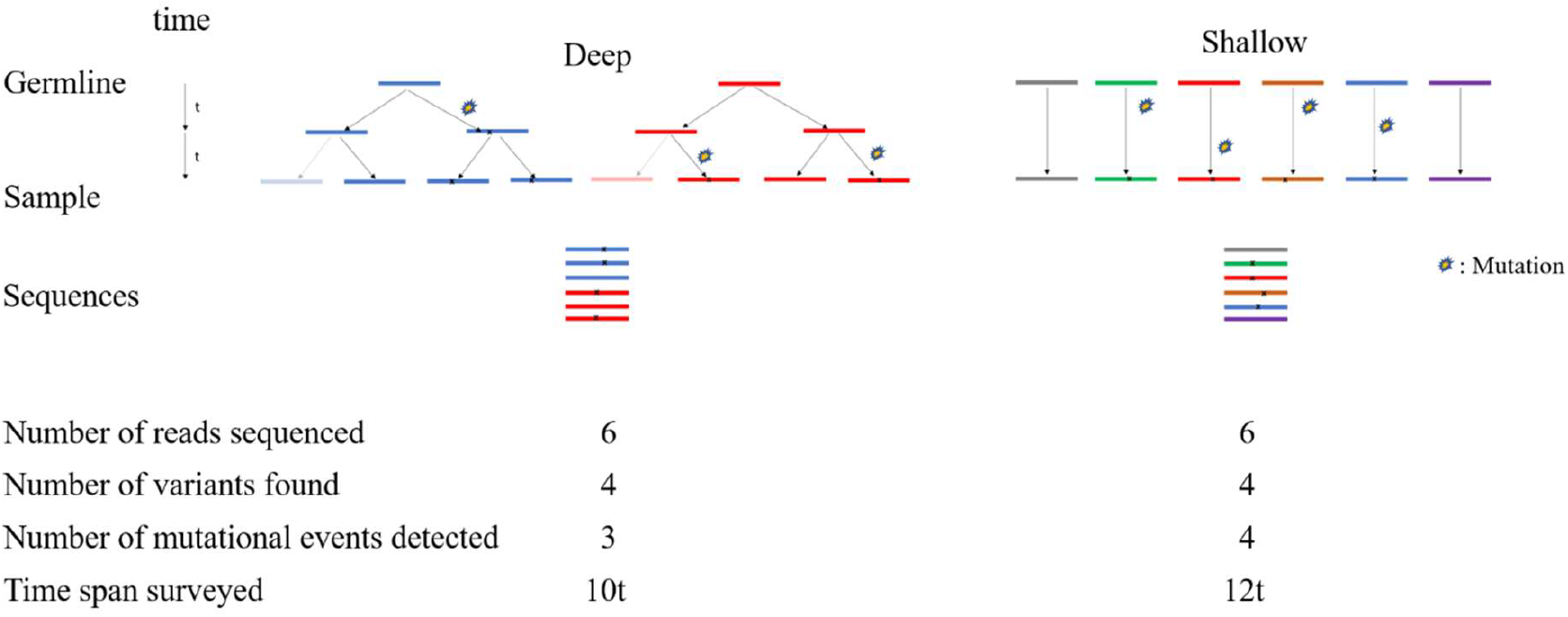
Using deep or shallow sequencing for detection of mutational patterns. Panel A: Illustrates the ancestral relationships between the sequences.

The figure highlights two potential disadvantages of using high compared to shallow sequencing depth for the same total amount of sequence data. The first is that the time span surveyed by the sequencing results is shorter using a deep sequencing approach. Therefore, the time mutagenic factors have to leave a characteristic imprint in the DNA is shorter (10 vs 12 time units in Figure 1). The second is the relationship between the number of mutational events and the number of variants present in the sequencing results. The higher the depth the more likely it is that a single mutational event is reflected by more than one read. In the figure, although in both protocols 6 reads were sequenced and 4 variants detected, only 3 mutational events were responsible for the 4 changes observed using the approach with higher depth. However, increasing depth allows a more accurate characterisation of the spectrum of changes that can be observed at a particular position.

In cases where only tumour material is available, the presence of germline variants can complicate detection of mismatch repair deficiency. In such cases it can be difficult to ascertain whether certain alleles have arisen through somatic mutations or were inherited from the parents. This has led to the use of “quasi monomorphic” microsatellites. These are microsatellites where germline variants are rare. Thus, the presence of variants is likely to reflect somatic mutations. Figure 2 shows the proportion of reads for a microsatellite (LR52, in Gallon et al 2020 ^10^) in a mismatch repair proficient and in a mismatch repair deficient sample.

**Figure 2:**
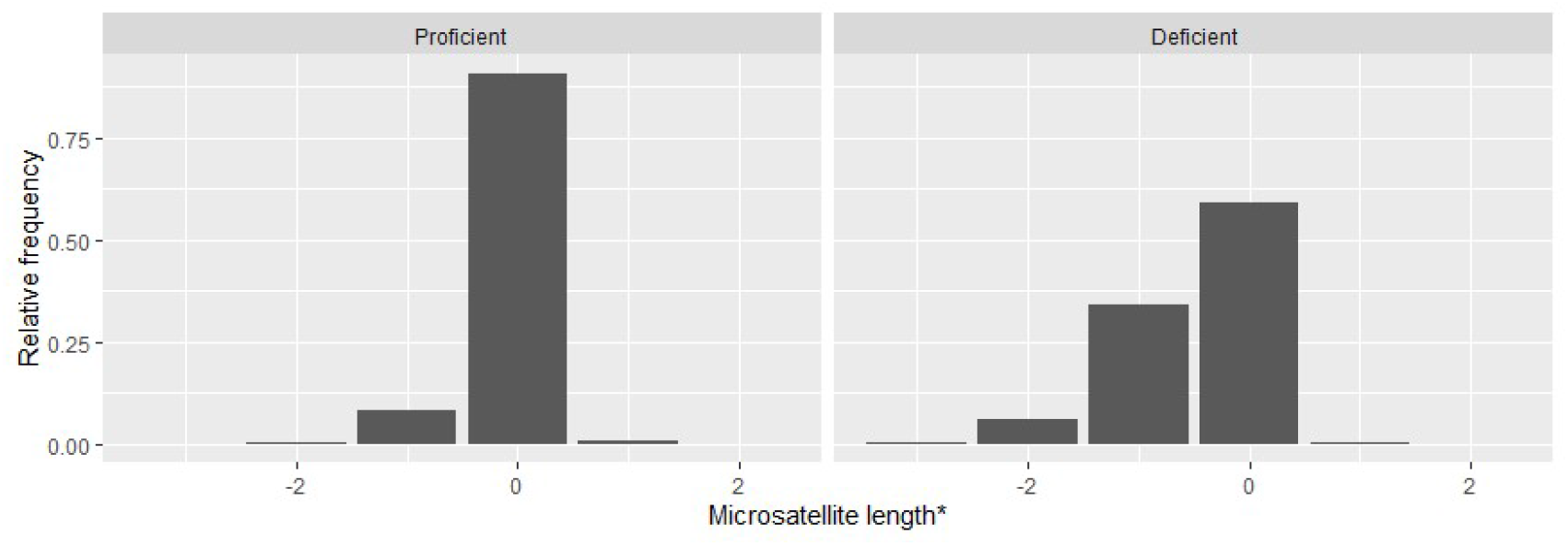
Distribution of reads in a MMR proficient and a MMR defficient sample. * Length difference with respect to the reference sequence (hg19)

**Figure 3.**
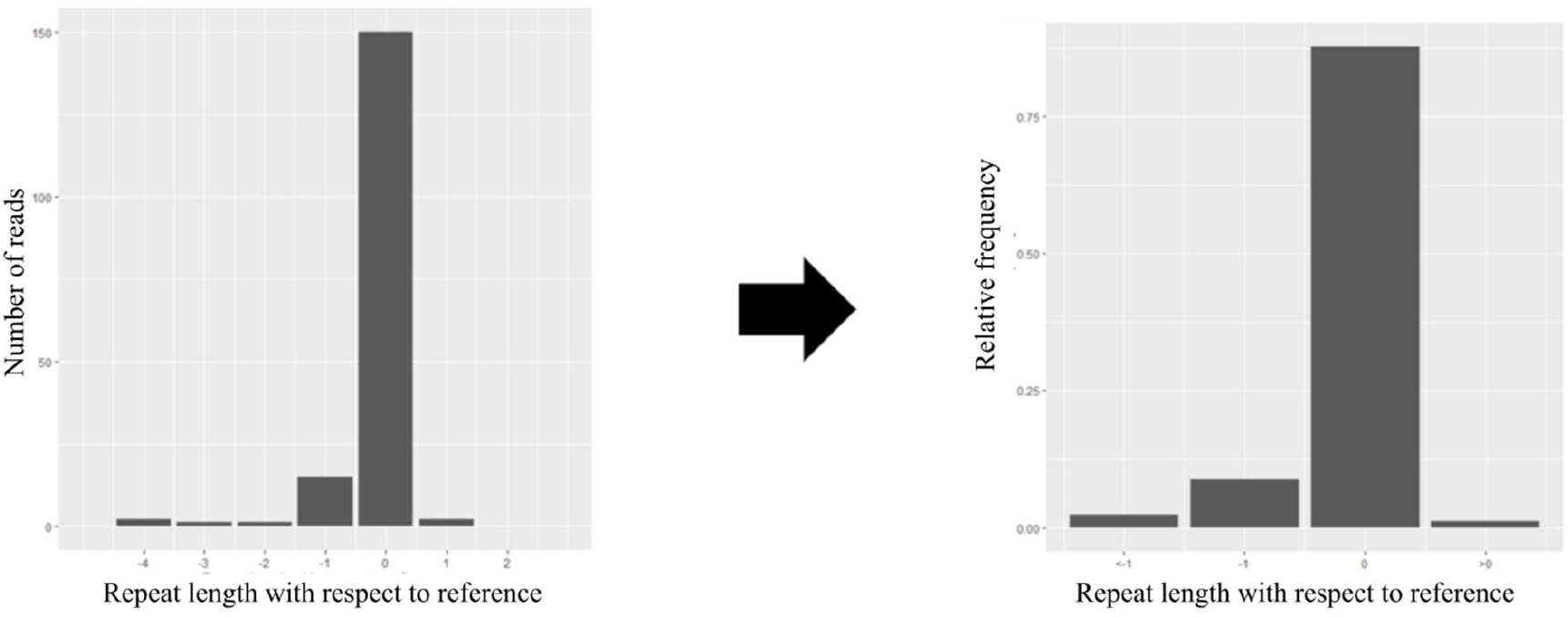
Binning alleles in a read group

**Figure 4:**
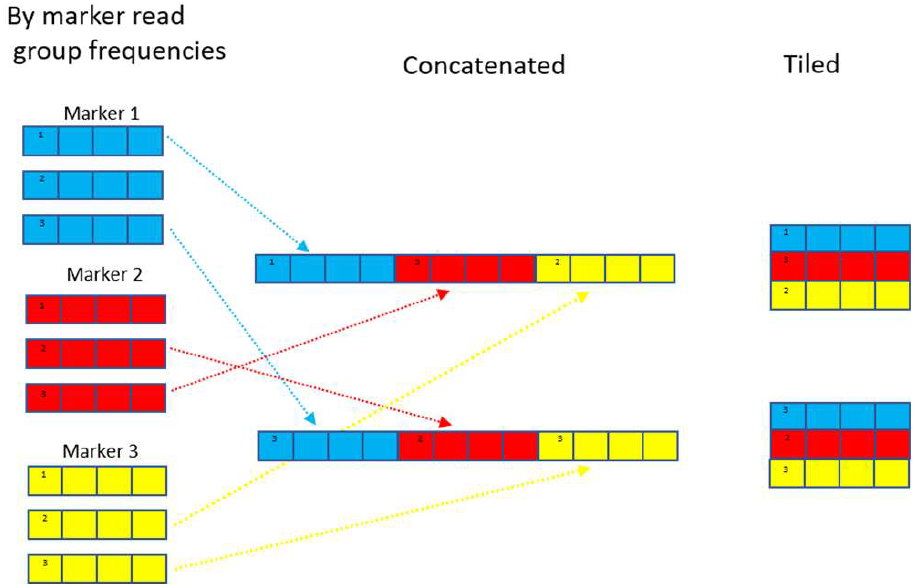
Data representations of marker combinations.

In the mismatch repair proficient sample, the sequencing detects reads representing deletions (e.g., lengths -1 and -2 in the figure) and insertions (length +1). The original publication ^11^ used unique molecular identifiers to conclude that such reads represent to a large degree amplification and sequencing artifacts. The MMR deficient sample has a higher frequency of non-reference length reads, in particular deletions, and a lower frequency of reference length reads (i.e., of length 0). The proportion of reference length reads has been often used to assess mismatch repair deficiency in tumours and non-malignant tissues (e.g. ^10,12^) and will be used here as a benchmark to judge other classification procedures. It is a summary measure that does not consider mutations of non-reference length alleles.

Here we use data from a set of samples from colorectal tumours whose mismatch repair status had previously been assessed using capillary electrophoresis to evaluate the performance of the different methods. Classification using alternative techniques, such as immunohistochemistry or sequencing of MMR genes, was not available. For colorectal tumours discordance between the results using different techniques has been reported to be in the order of 1-2% ^13-15^ and interpretation of our results should consider the accuracy of the original classification.

The aim of this report is to explore the influence of read depth on the ability to separate tumour sequence data according to the mismatch repair status of the original tumour using published microsatellite sequence data and artificial neural networks. We first use cross validation to explore the influence of read depth and the value of different models, and subsequently validate the results on an independent test set.

## 3. Material and methods

### 3.1 Data

We used published sequence data for short amplicons (150bp) targeting 24 short (between 7 and 12bp in length) mononucleotide repeats from MMR proficient and deficient paraffin embedded formalin fixed samples from colorectal tumours (EMBL-EBI European Nucleotide Archive, accession number PRJEB28394)^10^. Mismatch repair proficiency as ascertained using the MSI

Analysis System (Promega) is recorded. Samples classified as MSI-H were considered to be mismatch repair deficient and MSS or MSI-L samples as proficient. The data were split into two sets. A first set consists of data from 49 MMR proficient and 50 MMR deficient tumours and was used as training set in this paper. The second set includes data from 46 MMR proficient and 49 MMR deficient tumours and was only used as a test set (section 5.3). These sets were sequenced separately. The mean read depth in the training set was 6914 and 3853 in the test set.

### 3.2 Methods

Prior to analysis the reads were partitioned in groups of 12 and 200 reads. Additional read group sizes were considered in sections 4.1 and 4.3. Read groups were labelled according to the MMR proficiency of the sample of origin. For each read group MNR alleles were groups into four bins according to the deviation from the reference sequence length (deletions 2bp or longer, of 1bp, unchanged, and insertions) and their relative frequencies determined.

To assess the effect of using several markers, a read group was randomly selected and read group frequencies for the different markers were concatenated or tiled.

Here we present results obtained using simple artificial neural networks (ANN) implemented using tensorflow and keras through an R interface (https://tensorflow.rstudio.com/). We explored networks with up to five fully connected feed forward layers and, for marker combinations consisting of more than five markers, networks that included a single convolutional layer (see additional material for more details)

Unless stated otherwise, the results were obtained using four-fold cross validation, where selection of the validation set was done by sample, i.e., all the read groups belonging to a sample were either in the training or in the validation set.

The area under the curve (AUC) of the receiver operating characteristic curve was used as criterion for the performance of a classification method. We report AUCs by read group.

For single markers analysis was carried out using each of the 24 markers included in the original data. To explore the consequences of increasing the number of markers we generated random combinations of 2, 4, 6 and 8 markers. For each group size 25 combinations were randomly chosen (Listed in supplementary table 1).

Regularised logistic regression and a single node network were used to assess the performance of larger models. Additionally, for single markers, classification using the proportion of reference length reads was also included.

## 4 Results

### 4.1 Single marker analyses

We first explore classification for each of the 24 markers. Figure 5 represents the relationship between read group size and AUC using the reference length allele proportion as classification criterion.

**Figure 5:**
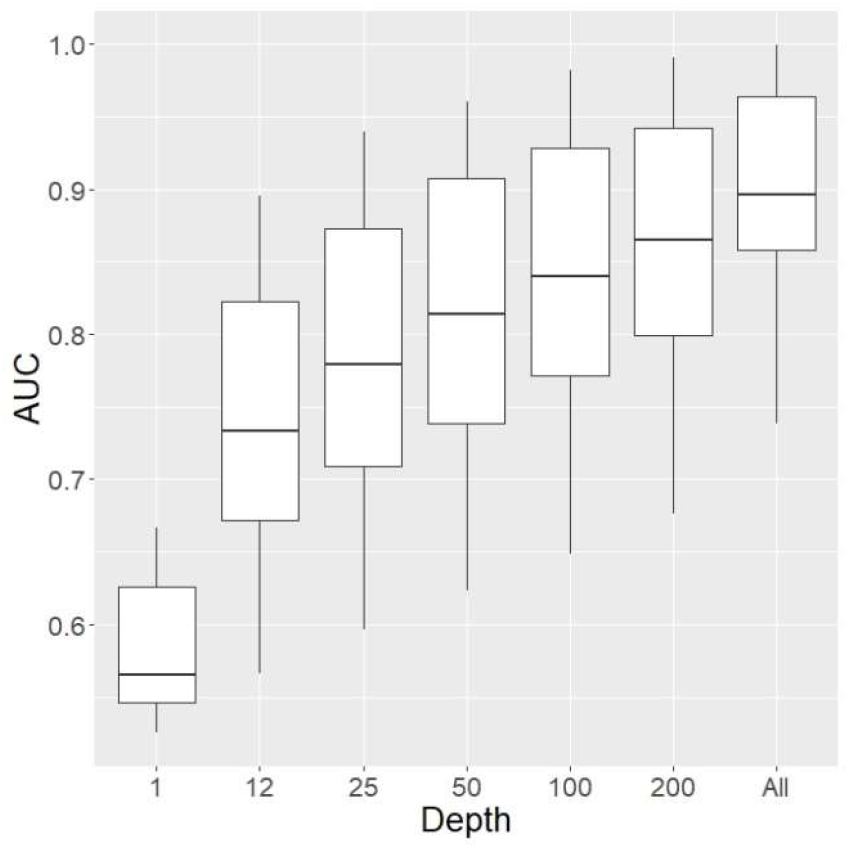
Classification using reference length read proportion. All: Using all available reads for each amplicon (median depth:6416)

As expected, the ability to classify improves with increasing read group size.

For read group sizes of 12 and 200 reads we investigate a range of models consisting of fully connected feed forward layers (Figure 6).

**Figure 6:**
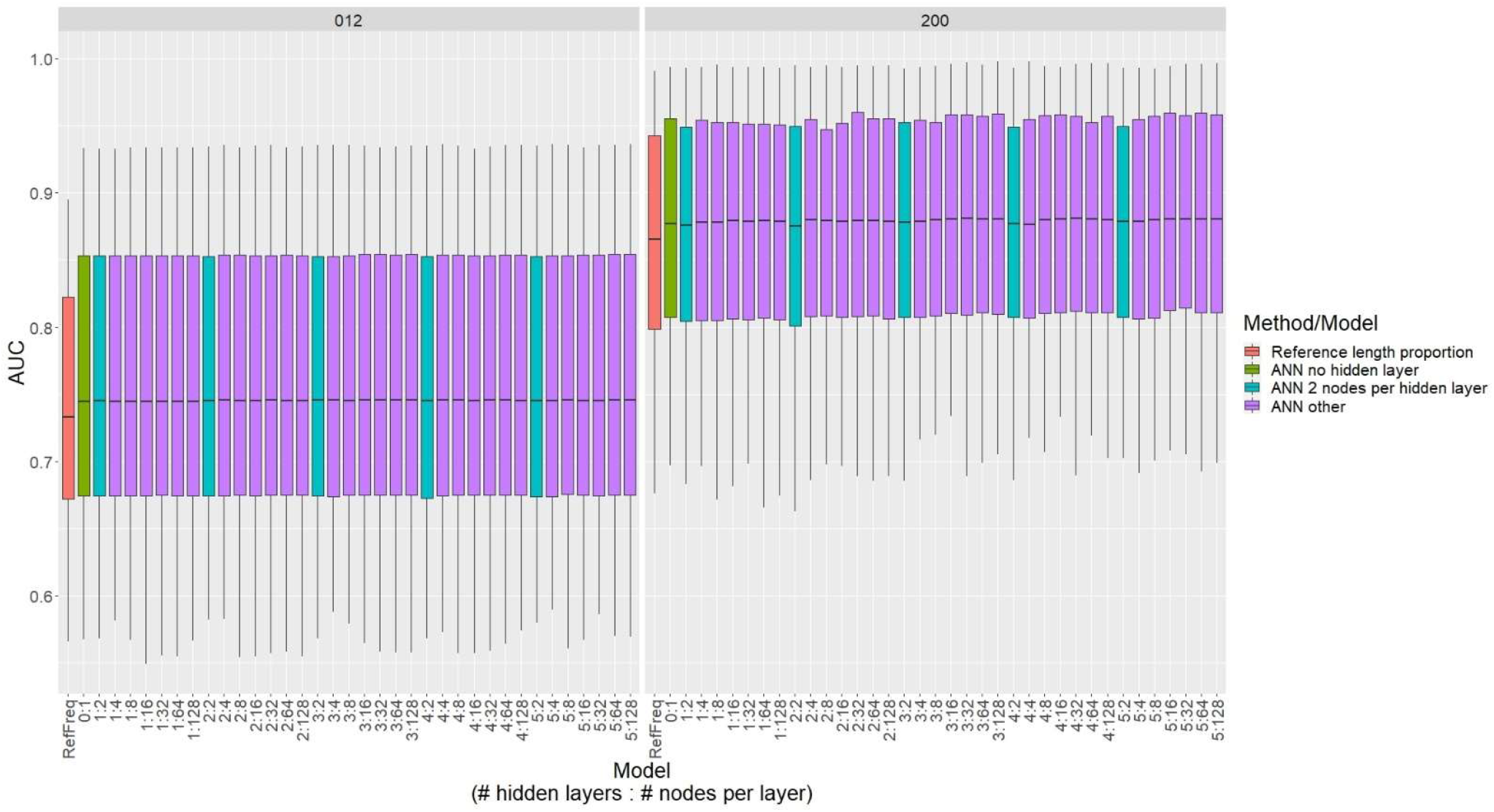
Single marker AUC (4-fold cross validation) for sequencing depths of 12 and 200.

Compared to the AUCs obtained using the reference length allele frequency, analyses of binned allele frequencies lead to an improvement (p<10^−5^ for any comparison). The results do not differ substantially across the different networks explored. At a depth of 200, 14 of the 24 markers achieve AUCs above 0.90 and 4 above 0.99 for any of the models.

### 4.2 Combinations of markers

The ability of groups of markers to separate MSI-H from MSS samples was explored for combination of 2,4,6 and 8 markers and investigated using 25 randomly chosen combinations (see methods) and 4-fold cross validation.

#### 4.2.1 Dense networks

The results are summarised in Figure 7. AUC increases with the number of markers and sequencing depth. It should be noted that the differences between the various networks is small (see supplementary figure S1).

**Figure 7:**
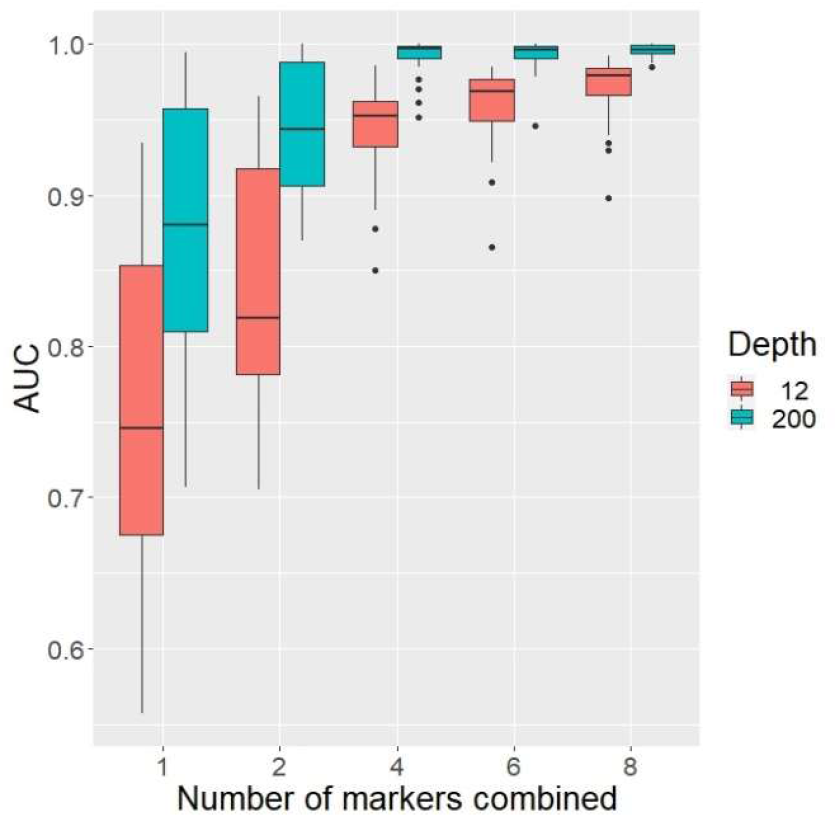
AUC for combinations of markers. 25 random combinations used for combinations of 2 to 8 markers and all 24 markers in the single marker analysis.

Only 6 combinations achieved an AUC above 0.99 at a read depth of 12. All of them consisted of 8 markers (see Fig 8). For a read depth of 200, 4 single markers (out of 24) and 74 (out of 100) combinations surpassed this threshold. Thus, we decided to concentrate on exploring a read depth of 12 in subsequent analyses.

**Figure 8:**
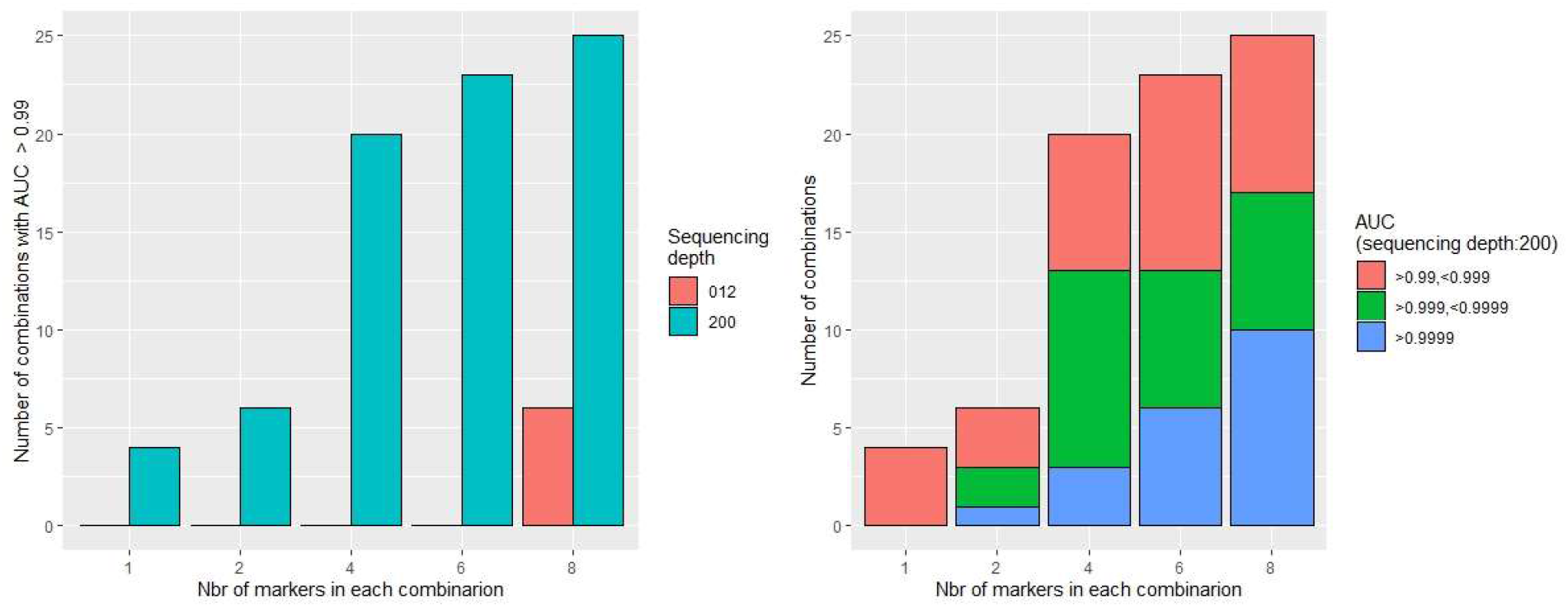
Number of combinations achieving AUCs above 0.99 for the different combinations. A: by sequencing depth, B: distribution for 200 fold coverage

#### 4.2.2 Use of a convolutional layer

The previous results suggest that variation in the number of layers or in the number of nodes per layer does not have a dramatical effect on discrimination performance. This is perhaps surprising. We thus explored an alternative representation of the input data by tiling the allele frequencies (see Figure 9 and methods Figure 4 in the methods section).

**Figure 9:**
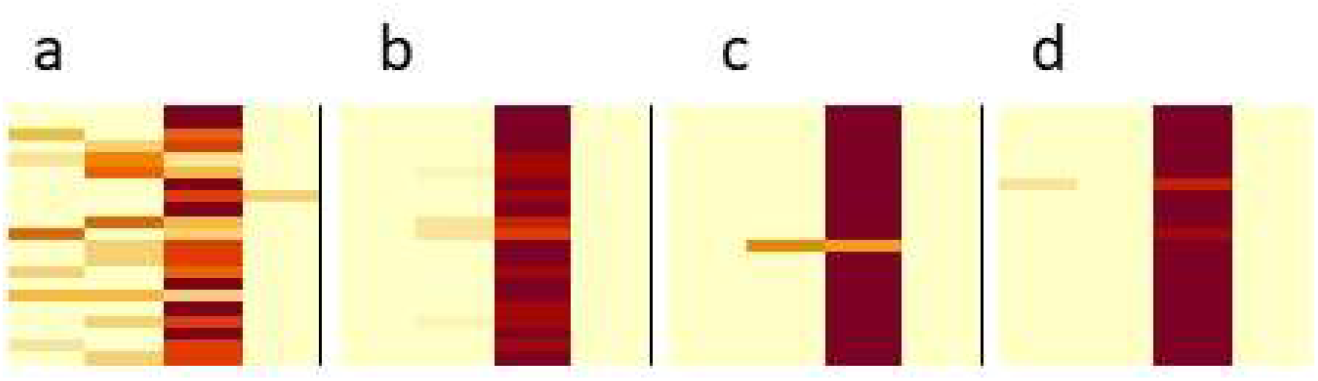
Representation of tiled data for two MSI-H (a and b) and two MSS (c and d) using a read depth of 200, all 24 markers (see methods).

Such a representation suggests that convolutional layers could be incorporated in the analysis. Here we only investigate the simplest models consisting of a single convolutional layer and the output layer using10-fold cross validation. The results for sequencing depth of 12 and for 6 and 8 markers are shown in Figure 10.

**Figure 10:**
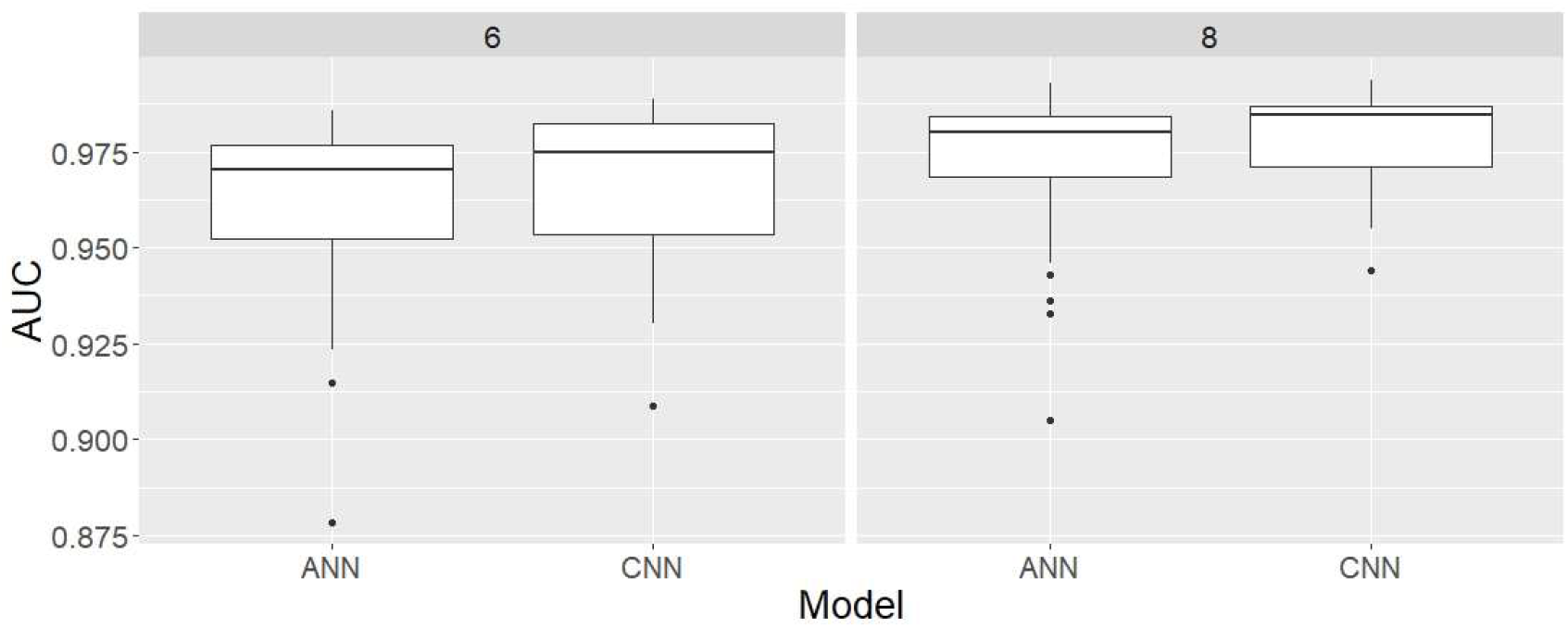
Comparing performance of the best dense network and a network containing one convolutional layer for a read depth of 12

Using a single convolutional layer significantly improves the performance in both sets (p=9×10^−8^ and p=3 10^−6^ for the sets of 6 and 8 marker combinations respectively, Wilcoxon signed-rank test).

### 4.3 Total number of reads

In practice, the interest will focus on allocating sequencing capacity. Figure 11 explores the relationship between the number of markers and classification ability when the total number of reads per sample is fixed. It suggests that, for a fixed total number of reads, higher AUCs are achieved by increasing the number of markers. However, with increasing number of markers the gain achieved by using additional markers is reduced.

**Figure 11:**
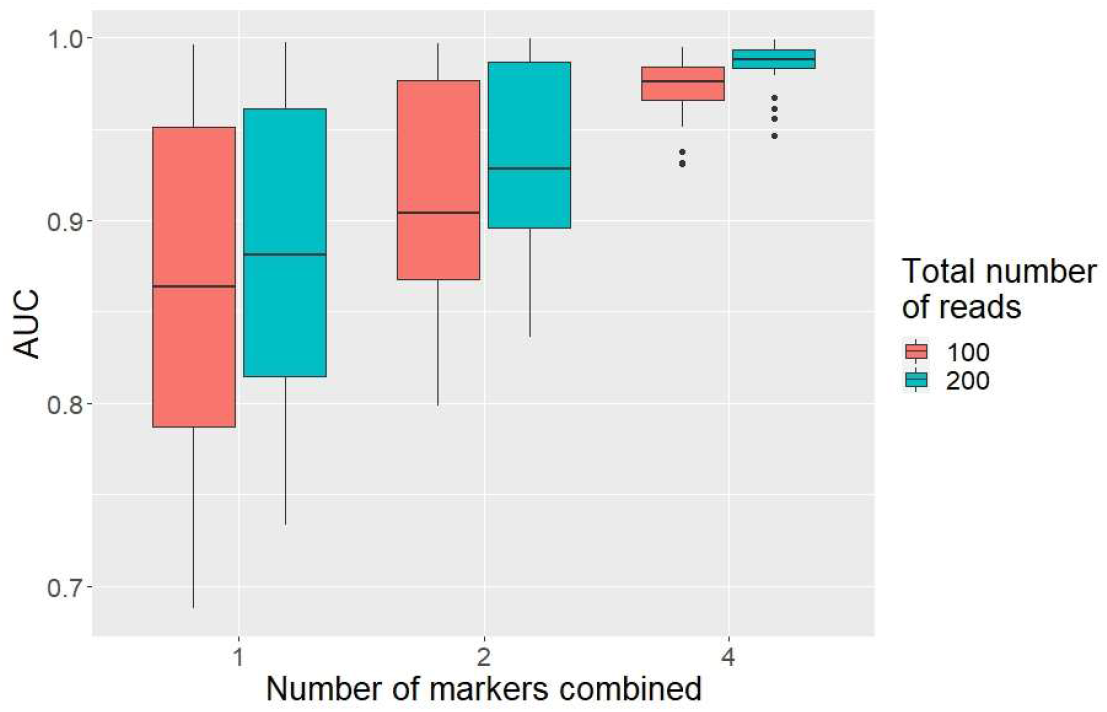
Distributing a fixed number of reads across varying number of markers

### 4.4 Classification of the test group

So far, our analyses have relied on cross validation using samples that were sequenced in one batch. In this section we examine the use of the training set to determine the parameters for each marker combination and evaluate their ability to classify a set of samples in the test set. To avoid optimising towards the test set we only investigated 50 combinations of two and four markers at a sequencing depth of 200. The 2 and 4-marker combinations were analysed using a model without a convolutional layer. The models were trained using 90% of the reads in the training group, the AUC determined for the remaining 10% and lack of improvement used as stopping criterion for the optimisation. For each combination the test group and the whole of the training group were reclassified using the parameters achieved at stopping time. The relationship between accuracy and AUC in the training and test sets for the 2 and 4 marker combinations is represented in Figure 12.

**Figure 12:**
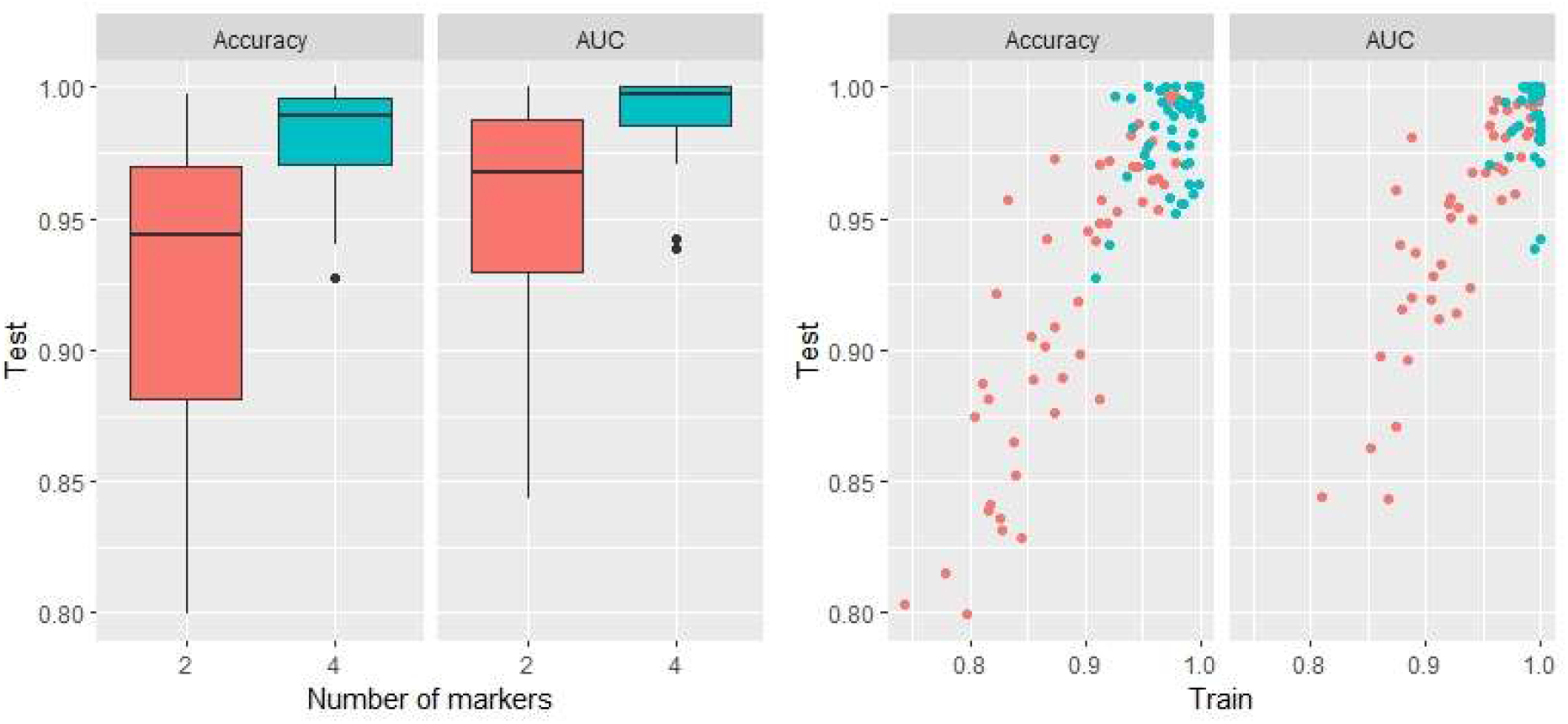
Performance of 50 2 and 4-marker combinations in training and test group at a read depth of 12

Among the 50 randomly chosen marker pairs median accuracy and AUC in the test set were 0.895 and 0.960 respectively; 5 combinations achieved an accuracy above 0.975 and 12 an AUC above 0.99. For the 4-marker combinations the median accuracy was 0.959 and the median AUC 0.999, 20 combinations achieved an accuracy above 0.975 and 18 an AUC above 0.99.

## 5. Discussion

Historically identification of microsatellite instability has focussed on counting the number of microsatellites that carry a mutation. The criteria for assessing whether a mutation has occurred at a particular microsatellite range from the emergence of novel modes in the allele length distribution to the reduction of the relative frequency of the alleles present in the germline normal tissues past a certain threshold. Often the distribution in non-cancerous tissues is used as a surrogate for the distribution in the germline. Here we used the relative frequencies of alleles as grouped by length directly. Underlying this approach is the assumption that for any microsatellite, mutations occur both in microsatellite stable and unstable tissues, and can be detected provided enough molecules are analysed, but that their frequency will differ in both groups and that this difference will eventually lead to differences in the allele length distributions. This circumvents the need of having to determine whether observed variants of a specific microsatellite are the result of *in vivo* mutations or of technical artifacts.

In the single marker analyses, comparison of the AUCs induced by using the frequency of the putative germline allele as classification criterion and those obtained using the frequencies of all bins indicate that additional information can be obtained by considering the extent of the shift in microsatellite length (see section 4.1 and Fig 6). This could reflect the underlying mutational mechanisms or recurrence of mutations in already mutated alleles and could be of interest in examining tumours where, during tumour evolution, the population of MMR deficient cells has either remained constant or grown slowly over prolonged periods and may still represent a modest proportion of the sample DNA.

We used ANNs for the analysis. This choice was mainly motivated by their flexibility, the potential to harness interactions between input features and possibility of knowledge transfer. Our data suggest that appropriate training sets can be obtained by splitting data from samples sequenced at a high depth and combining data from different markers from the same sample. The results using a convolutional layer suggest that multi-marker data can be represented as two dimensional arrays and analysed in a way similar to that used for images. However, in this context it should also be considered that the requirement of classification invariance with respect to rotation and translation of the whole image are of little interest when using monomorphic markers.

Different markers or marker combinations vary in their sensitivity to MMR deficiency. The work presented here aimed to explore the impact of sequencing depth and number of sites on MMR detection ability and not to identify optimal sets of markers. We used relatively short microsatellites and summarised the length distribution by condensing the allele sizes into 4 bins. Longer microsatellites are used in many assays (e.g., ^16,17^). They tend to be more sensitive to MMR deficiency and accuracies above 97% have been described for single markers analysed by fragment length analysis ^17^. Their use in sequence-based assays has been hampered by the error rate of available sequencing technologies and their germline variability can limit their value when no matching normal tissue is available. For such microsatellites both artefactual and *in vivo* changes can involve a larger number of repeat units and increasing number of bins could be advantageous.

Using random combinations of markers, we were able to obtain reliable classification at relatively low sequencing depth of 12. This contrasts with the findings from Cortes-Ciriano et al. (2017) ^18^ who investigated whole genome and whole exome sequencing data, concluded that sequencing depths below 30 impair the ability to detect microsatellite instability. Other targeted sequencing-based procedure used a larger number of reads ranging from 5 to 5000 (for a recent review see ^19^). The numbers of markers in such assays varied with up to several thousand markers scored for methods using whole genome sequencing, while assays relying on targeted enrichment used panels from 2 to up to several thousand microsatellites ^19^. However, the latter used sequencing depths of several thousand reads. Gallon et al. 2020 ^10^ proposed a subset of 6 markers that achieved an accuracy of around 0.96 when consensus sequences for 25 different unique molecular identifiers were used (the average number of reads required to capture the 25 identifiers was not shown). Even if 25 reads per marker were used that is 50% more reads than 12 reads per marker for 8 markers. Both the small number of markers and the low read depth are of interest since they allow to reduce the amount of DNA required for each sample and to increase the number of samples assessed in a sequencing run.

Requiring a low sequencing depth is also of interest in the context of shallow whole genome sequencing. We did not explore sequencing depths of 1 or 2. Such low depths could be of interest in the context of single cell sequencing. Using cross validation and all 24 markers we reached an AUC of 0.91 at a sequencing depth of 1 and 0.95 for a depth of 2 (data not shown). However, these values are likely to underestimate classification performance that could be obtained in single cell analyses since the MMR deficient tumour samples will contain MMR proficient material from contaminating normal material or from infiltrating circulating cells. When data for different markers are combined, reads from MMR deficient and proficient cells will be mixed. The limit of detection for the Promega Oncomate MSI assay used here as reference method, is reported to be 20% or lower ^20^. Thus a sample containing 30% MMR deficient cells would be classified as MMR deficient, but at a read depth of 1, 3% of all 10 markers combinations would contain no sequences from the MMR deficient cells and more than half of the makers will be represented by data from MMR proficient cells in 97% of 24 marker combinations. This is not an issue when single cell data are analysed.

Although the widespread introduction of whole genome sequencing raises the question of the value of targeted amplification-based methods in general, currently the latter are still likely to be cheaper and more suitable for screening large numbers of samples. More generally, increased microsatellite instability as a functional correlate for MMR deficiency can be helpful in situation where sequencing has failed to detect defects in mismatch repair genes or when the functional significance of particular variants is unclear. In the context of whole genome sequencing, our approach could be useful for exploring differences in the response of particular regions of the genome to mismatch repair deficiency.

In summary, our results demonstrate the use of artificial neural networks to detect mismatch repair deficiency. They indicate that sequencing depth can compensate for target number. However, they also suggest that for a fixed total number of reads per sample increasing sensitivity by increasing the number of targets is more efficient than by increasing per target sequencing depth.

## Additional material

### Hyperparameters

For hidden layers ReLu was used as activation function and binary cross entropy for the output layer. Adam was used as optimiser and with exception of the learning rate default parameters were retained. For cross-validation, lack of improvement of the AUC in the validation group was used as stopping and the following parameters gids were used: Stopping criterion: For fully connected feed forward networks: Learning rate: 0.1,0.01,0.001,0.0001; Number of hidden layers: 0-5; Number of nodes per layer 0,1,2,4,8,16,32,64,128; maximum numbers of epochs:200. For networks with a convolutional layer: filter dimensions: 2×2, 3×3 and 4×4; filter number: 2,4,8,16,32,64,128,256; Learning rate: 1e-1,1e-2,1e-4,1e-5,1e-6; maximum numbers of epochs: 80. No padding.

### Marker combinations

Using the nomenclature from Gallon et al. 2020^10^ the following marker combinations were used:

Single markers: DEPDC2, GM01, GM07, GM09, GM11, GM14, GM17, GM22, GM26, GM29, IM16, IM49, LR10, LR11, LR17, LR20, LR24, LR36, LR40, LR44, LR46, LR48, LR49, LR52.

2 marker combinations: DEPDC2.GM26, GM07.GM17, GM07.GM29, GM07.IM16, GM07.LR44, GM11.GM26, GM17.GM26, GM22.GM26, GM22.LR11, GM22.LR40, GM26.LR10, GM26.LR49, IM16.LR36, IM16.LR48, LR10.LR24, LR10.LR40, LR11.LR36, LR11.LR46, LR17.LR52, LR24.LR46, LR36.LR49, LR40.LR46, LR40.LR48, LR44.LR46, LR46.LR49

4 marker combinations: DEPDC2.GM09.IM16.LR24, DEPDC2.GM14.LR24.LR40, DEPDC2.GM17.LR11.LR52, GM01.GM09.IM49.LR44, GM01.GM11.GM14.LR44, GM01.GM22.IM49.LR52, GM01.LR17.LR24.LR36, GM07.GM09.LR10.LR36, GM07.GM11.IM16.LR10, GM07.GM17.LR10.LR20, GM07.GM26.LR10.LR11, GM07.LR17.LR24.LR52, GM09.GM11.GM14.LR17, GM09.GM14.GM17.LR52, GM11.GM14.GM17.LR48, GM11.GM29.LR17.LR49, GM14.GM17.LR11.LR20, GM17.GM29.LR17.LR24, GM22.GM26.LR49.LR52, GM22.IM49.LR10.LR44, GM26.GM29.LR24.LR44, GM26.IM16.LR24.LR46, IM16.IM49.LR36.LR52, IM16.LR10.LR46.LR48, LR10.LR20.LR24.LR48

6 marker combinations: DEPDC2.GM01.GM07.GM09.LR44.LR48, DEPDC2.GM01.GM09.LR11.LR20.LR46, DEPDC2.GM07.GM09.GM11.LR24.LR48, DEPDC2.GM07.GM17.LR17.LR24.LR52, DEPDC2.GM09.IM49.LR10.LR20.LR46, DEPDC2.GM11.GM17.LR10.LR20.LR24, DEPDC2.GM14.LR10.LR46.LR49.LR52, DEPDC2.GM29.IM16.LR11.LR46.LR49, DEPDC2.LR10.LR24.LR36.LR48.LR49, GM01.GM07.GM11.GM17.GM29.LR44, GM01.GM11.LR40.LR44.LR48.LR52, GM01.GM17.GM26.LR17.LR20.LR40, GM01.GM22.LR11.LR17.LR44.LR48, GM01.GM26.LR36.LR48.LR49.LR52, GM01.IM16.LR17.LR20.LR44.LR49, GM07.GM09.GM14.IM49.LR24.LR48, GM07.GM17.LR10.LR20.LR36.LR44, GM09.GM26.IM16.LR10.LR24.LR36, GM09.IM49.LR36.LR44.LR48.LR49, GM11.GM14.GM29.LR11.LR40.LR46, GM17.LR11.LR24.LR46.LR48.LR52, GM22.GM29.IM16.LR10.LR48.LR49, GM22.GM29.IM16.LR40.LR44.LR46, GM26.LR10.LR17.LR46.LR49.LR52, IM16.LR20.LR36.LR40.LR46.LR4

8 marker combinations: DEPDC2.GM01.GM07.GM09.GM17.IM16.LR36.LR40, DEPDC2.GM01.GM09.GM14.GM29.LR24.LR40.LR49, DEPDC2.GM07.GM09.GM14.GM22.IM49.LR40.LR48, DEPDC2.GM07.GM11.GM22.IM49.LR17.LR46.LR52, DEPDC2.GM07.GM22.IM49.LR20.LR40.LR44.LR49, DEPDC2.GM14.GM17.GM26.LR20.LR24.LR46.LR49, DEPDC2.GM17.GM22.GM26.GM29.LR36.LR46.LR52, GM01.GM07.GM14.GM17.LR10.LR20.LR24.LR36, GM01.GM09.GM14.GM26.GM29.IM49.LR36.LR46, GM01.GM09.GM22.GM26.LR11.LR17.LR20.LR36, GM01.GM11.GM17.GM22.LR17.LR20.LR24.LR40, GM01.GM11.GM26.GM29.IM16.LR11.LR46.LR48, GM01.GM14.GM22.IM16.LR11.LR17.LR40.LR44, GM07.GM09.GM26.GM29.LR20.LR36.LR46.LR48, GM07.GM11.GM14.GM17.GM26.LR24.LR44.LR48, GM09.GM14.LR10.LR20.LR36.LR40.LR49.LR52, GM09.GM17.IM49.LR24.LR40.LR44.LR49.LR52, GM11.GM14.GM29.IM49.LR17.LR36.LR40.LR46, GM11.LR17.LR20.LR24.LR40.LR46.LR49.LR52, GM14.GM29.IM16.IM49.LR10.LR20.LR46.LR48, GM14.IM49.LR11.LR17.LR36.LR40.LR49.LR52, GM22.IM49.LR11.LR17.LR20.LR24.LR44.LR49, GM26.LR11.LR17.LR24.LR36.LR44.LR48.LR49, GM29.LR11.LR20.LR24.LR36.LR40.LR46.LR49, IM16.IM49.LR10.LR11.LR17.LR44.LR48.LR52

## Supplementary Figures

**S1:**
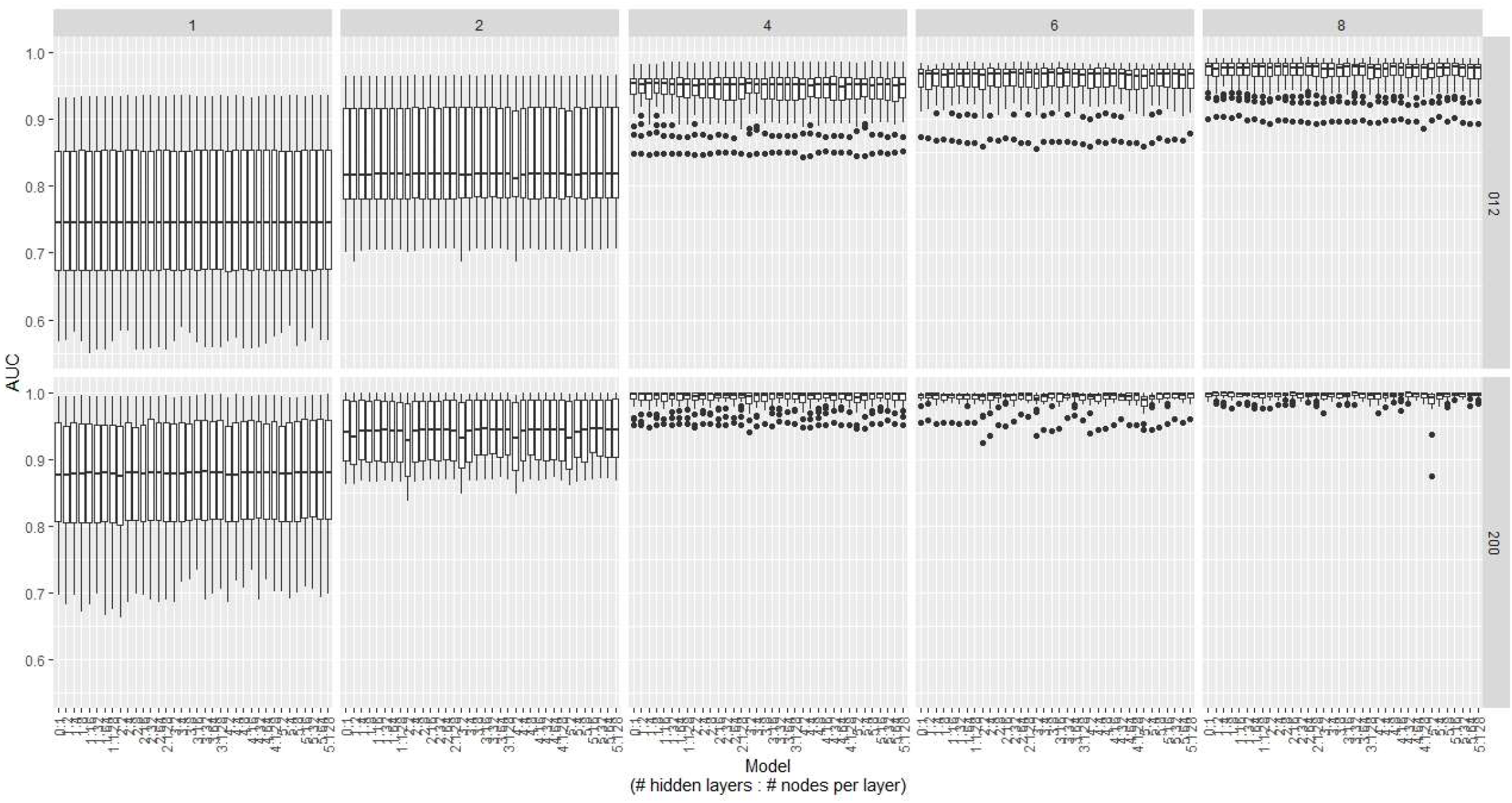

## Notes

### Competing Interest Statement

Richard Gallon,John Burn, Michael Jackson and Mauro Santibanez Koref: named inventors on patents covering the microsatellite instability markers analyzed: WO/2018/037231 (published March 1, 2018), WO/2021/019197 (published February 4, 2021), and GB2114136.1 (filed October 1, 2021).

